# Transcriptomic landscape of early hair follicle and epidermal development

**DOI:** 10.1101/2022.11.03.515012

**Authors:** Ana-Marija Sulic, Rishi Das Roy, Verdiana Papagno, Qiang Lan, Riikka Saikkonen, Jukka Jernvall, Irma Thesleff, Marja L Mikkola

## Abstract

Morphogenesis of ectodermal organs, such as hair, tooth, and mammary gland, starts with the formation of local epithelial thickenings, or placodes, but it remains to be determined how distinct cell types and differentiation programs are established during ontogeny. Here, we use bulk and single-cell transcriptomics and pseudotime modelling to address these questions in developing hair follicles and epidermis, and produce a comprehensive transcriptomic profile of cellular populations in the hair placode and interplacodal epithelium. We report previously unknown cell populations and marker genes, including early suprabasal and genuine interfollicular basal markers, and propose the identity of suprabasal progenitors. By uncovering four different hair placode cell populations organized in three spatially distinct areas, with fine gene expression gradients between them, we posit early establishment of cell fates. This work is accompanied by a readily accessible online tool to stimulate further research on skin appendages and their progenitors.

## INTRODUCTION

Skin and its appendages provide a protective barrier that has essential thermoregulatory, sensory and metabolic functions. Mammalian skin is composed of the outermost epidermis and the dermis, separated by a basement membrane (Fuchs, 2007). Skin appendages include hair, teeth, mammary, salivary and sweat glands, feathers in birds and scales in reptiles and fish. Although these organs differ in their adult form and function, their early development shares multiple common features at both morphogenetic and molecular levels (Mikkola and Millar, 2006; Biggs and Mikkola, 2014). Hair follicle (HF) is one of the best-studied models of skin appendage development. It is easily accessible on the surface of the embryo, its development proceeds via well-defined morphological stages, and a large repertoire of tools is available for its genetic manipulation in mice. Similar to other skin appendages, HF development begins by the formation of a local epithelial thickening, a placode (Pispa and Thesleff, 2003). Placode cells give rise to at least eight molecularly distinct cell types in the fully formed HF, as well as stem cells (SC) that fuel the cyclic regeneration of HFs (Levy et al., 2005; Saxena et al., 2019). Differential expression of placode markers (Heath et al., 2009) and the stratified structure of the placode (Ahtiainen et al., 2014) indicate that cellular heterogeneity exists early on. Yet, the degree and composition of this heterogeneity, and the time of establishing cell fates, remains incompletely understood. With emergence of single cell technologies such as single-cell RNA sequencing (scRNA-seq), it is now possible to dive deep into the cellular and molecular heterogeneity of different organs and explore their cellular composition, transcriptomic landscape, and differentiation trajectories, giving us an unparalleled look at ontogeny.

The adult epidermis is a constantly renewing stratified tissue, consisting of the proliferative basal, and differentiated spinous, granular, and cornified layers of keratinocytes, each with characteristic molecular profiles and morphological features (Moreci and Lechler, 2020). Apart from keratinocytes, the primary cell type found in the skin epidermis, it also houses melanocytes, light-touch sensing Merkel cells, and Langerhans cells, the tissue-resident macrophages (Fuchs, 2007). The specification of the epidermal fate takes place soon after gastrulation and relies on the induction of the epidermal lineage-specific transcription factor p63 in the embryonic surface epithelium (Mills et al., 1999; Yang et al., 1999). The expression of the simple epithelia markers Krt8/Krt18 are replaced by the expression of Krt5/Krt14 at embryonic day (E) 9.5 in the mouse (Byrne et al., 1994). The first stratification of the single layer of multipotent epithelial cells takes place around E11.5 with the appearance of the periderm, a layer of flat cells that cover and protect the developing epithelium until late embryogenesis (Hammond et al., 2019). Soon after, intermediate suprabasal cells emerge, switch on expression of Krt1/Krt10 and later mature into the spinous layer. As development proceeds, spinous cells move upwards, give rise to granular cells that gradually compact and lose their cell organelles, forming the outmost stratum corneum, where the keratinization process is completed (Moreci and Lechler, 2020). All these steps of epidermal differentiation rely on multiple transcriptional regulators and signaling pathways, the canonical Notch signaling pathway playing a central role (Rangarajan et al., 2001; Blanpain et al., 2006; Nowell and Radtke, 2013).

Concomitant with epidermal morphogenesis, HF development begins during embryogenesis, and in mice is completed within the first two postnatal weeks (Sennett and Rendl, 2012; Biggs and Mikkola, 2014). Murine HFs are induced in three successive waves as a result of an unknown inductive signal(s) thought to arise in the mesenchyme. Subsequently, a regular array of hair follicle primordia emerges, the first (a.k.a. primary) ones at late E13.5, maturing into placodes by E14.5 (Sennett and Rendl, 2012; Biggs and Mikkola, 2014). Placode morphogenesis is associated with cell cycle cessation and is driven by intercalation and centripetal migration of placode-committed HF progenitors (Ahtiainen et al., 2014; Cetera et al., 2018). Accordingly, actin-mediated cell rearrangements are necessary for early HF morphogenesis (Ahtiainen et al., 2014; Cetera et al., 2018). Also, HF stem cells (HFSCs) are specified early during hair follicle morphogenesis (Nowak et al., 2008; Xu et al., 2015; Ouspenskaia et al., 2016). Nascent placodes signal to the adjacent mesenchymal fibroblasts to form the dermal condensate (DC), precursor of the mature dermal papilla, which is essential for continuing HF development and maintenance (Sennett and Rendl, 2012; Huh et al., 2013). Placode-DC crosstalk stimulates the downgrowth of the epithelium, forming the hair germ at E15 (Biggs and Mikkola, 2014). Thereafter, the follicular epithelium extends further into the dermis, engulfs the dermal papilla, and differentiation of the multiple cell types that constitute the mature HF begins (Sennett and Rendl, 2012).

Molecular regulation of placode formation has been extensively studied during past two decades. The canonical Wnt/β-catenin pathway is at the helm of HF induction: deletion of β-catenin in the developing epidermis results in the absence of all signs of hair placode formation (Huelsken et al., 2001; Zhang et al., 2009), while its forced activation induces precocious hair placode and programs the entire skin to HF fate (Närhi et al., 2008; Zhang et al., 2009). Placode maintenance, in turn, relies on the Eda/Edar/NF-κB pathway: in the absence of Eda signaling, primary hair follicle morphogenesis halts at a pre-placode stage (Schmidt-Ullrich et al., 2006; Fliniaux et al., 2008), whereas overexpression of *Eda* drives the formation of enlarged placodes (Mustonen et al., 2004; Ahtiainen et al., 2014). In contrast to Wnt and Eda pathways, the Bmp signaling inhibits placode formation and the noggin-mediated suppression of Bmp signaling in epithelium is a prerequisite for hair placode morphogenesis to take place (Botchkarev et al., 1999; Mou et al., 2006; Pummila et al., 2007). Sonic hedgehog (Shh) pathway is not necessary for placode formation, but is rather required for HF morphogenesis to proceed beyond the early germ stage and for DC formation (St-Jacques et al., 1998; Chiang et al., 1999; Qu et al., 2022). It plays an essential role in HFSCs specification: an antagonist interplay between Wnt/β-catenin and Shh signaling activity during early HF morphogenesis specifies the prospective stem cells (Xu et al., 2015; Ouspenskaia et al., 2016).

In this study, we examine cellular heterogeneity of the hair placode in the context of the developing skin, with the specific aim to obtain a comprehensive readout of the transcriptional topography of early hair development and interfollicular epithelium (IFE). Although efforts have been made in elucidating the transcriptional profile of the hair placode, they have relied on averaged transcriptomes provided by bulk RNA-seq and microarrays (Sennett et al., 2015; Tomann et al., 2016), whereas recent scRNA-seq studies mainly concentrated on the dermal niche (Gupta et al., 2019; Qu et al., 2022), or HFSCs of whisker follicles (Morita et al., 2021). Here, we combine in-depth transcriptional analysis of mature hair placodes using both bulk and scRNA-seq methods. Our study revealed placode and developing interfollicular epithelium gene signatures, as well as different spatially and genetically distinct cell populations constituting hair placode, and fine gene expression gradients between them, suggesting cell fates are already established at the hair placode stage. Cluster analysis and pseudotime modelling allowed discovery of early markers of suprabasal cells and genuine IFE basal markers. In addition, we provide a comprehensive comparison of bulk and scRNA-seq datasets and explore the advantages and disadvantages of both approaches. All of our bulk and scRNA-seq data are shared as a resource in an easily searchable online database.

## RESULTS

### Bulk RNA-sequencing identifies established and novel hair placode markers

To determine comprehensive hair placode and IFE gene signatures we profiled E14.5 mouse back skin epithelium using RNA-seq. Dorso-lateral skin of E14.5 embryos was micro-dissected and epithelium and mesenchyme were separated, followed by dissociation of the epithelial cells to obtain single cell suspension **(Figure 1A)**. To enrich the sample for placode cells we took advantage of the cell surface marker mucosal vascular addressin cell adhesion molecule 1 (MAdCAM-1) that has been previously shown to be expressed in hair placodes (Nishioka et al., 2002). Whole mount 3D confocal imaging confirmed focal MAdCAM-1 expression in the skin epithelium in a pattern that matched *Fgf20-LacZ* allele, a known early hair placode marker (Huh et al., 2013) **(Figure 1B)**. We used fluorescence-activated cell sorting (FACS) to obtain MAdCAM-1 positive (placode-enriched) and MAdCAM-1 negative (IFE) cells **(Figure S1A)** and performed RNA sequencing of seven biological replicates. On average, RNA-seq produced 47 million aligned reads per sample. We detected 26 007 genes, of which 5 502 were significantly differentially expressed (adjusted p-value <0.05) between placode-enriched (hereafter placode) and IFE cell populations; of these 3 115 had fold difference 1.5x or higher **(Figure S1B, Table S1)**.

**Figure 1.**
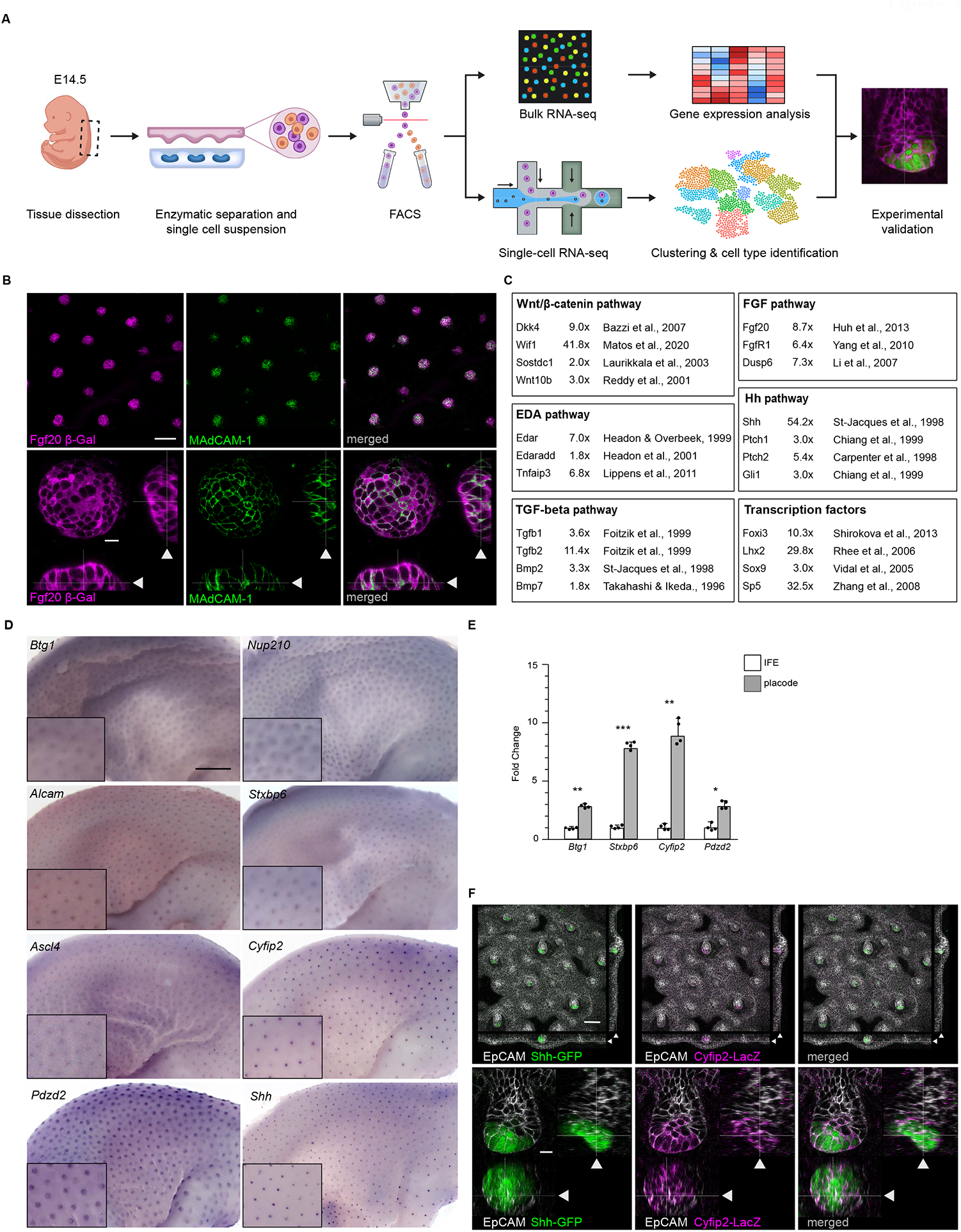
Identification of hair placode markers by bulk RNA-seq. (A) Experimental pipeline used for transcriptional profiling of hair placodes and interfollicular epithelium. (B) Optical sections (planar and sagittal views; arrowhead indicates section plane) of confocal microscopy images from *Fgf20*^*+/LacZ*^ mouse back skin (E14.5) labelled with antibodies against MAdCAM-1 (green) and β-galactosidase (magenta) to visualize hair placodes. Scale bar, 100 μm (upper panels) and 10 μm (lower panels). (C) Established hair placode markers obtained from RNA-seq of MAdCAM-1 positive mouse back skin epithelial cells (E14.5). Representative marker genes (Fc >1.5x), and the relevant references, are listed for each pathway. (D) Whole mount in situ hybridization (WMISH) using probes specific for placode markers *Btg1, Nup210, Alcam, Stxbp6, Ascl4, Cyfip2, Pdzd2*, and *Shh* (n = 5). Scale bar, 1 mm. (E) qRT-PCR on replicate samples for *Btg1, Stxbp6, Cyfip2* and *Pdzd2* (n = 4). Significance was assessed with two tailed Student’s T-test, *p < 0.05; **p < 0.01; ***p < 0.001. Error bars represent SD. (F) Optical sections (planar and sagittal views; arrowhead indicates section plane) of confocal microscopy images from *Cyfip2*^*+/LacZ*^*;Shh*^*+/GFPCre*^ E15.5 mouse back skin labelled with antibodies against EpCAM (white), β-galactosidase (magenta) and GFP (green) to visualize hair germs. Scale bar, 100 μm (upper panels) and 10 μm (lower panels).

Hierarchical clustering of differentially expressed genes showed placode and IFE biological replicates clustering together **(Figure S1C)**. Comparison of our data with established hair placode marker genes from the literature validated our sorting strategy. Placode population was, among others, defined by members of Wnt/β-catenin pathway (*Dkk4, Wif1, Sostdc1, Wnt10b*), Eda-pathway (*Edar, Edaradd, Tnfaip3*), TGF-β pathway (*Tgfb1, Tgfb2, Bmp2, Bmp7*), Fgf pathway (*Fgf20, FgfR1, Dusp6*), and the hedgehog pathway (*Shh, Ptch1, Ptch2, Gli1*), as well as transcription factors (TFs) *Foxi3, Lhx2, Sox9*, and *Sp5* **(Figure 1C, Table S1)**. Apart from high enrichment of genes involved in key signaling pathways, by means of Gene Set Enrichment Analysis (GSEA) (Mootha et al., 2003; Subramanian et al., 2005) we could also see enrichment of genes involved in substrate dependent cell migration (*Slit2, Robo1, Ntn1, Nrp1, Sema3a*), regulation of hair cycle (*Tnf, Tgfb2, Krt17, Wnt10b*), chemokine signaling (*Cxcl14, Cxcl9, Ccr10*), and glycan biosynthesis (*St6gal1, Mgat3, Man1a*) **(Figure S1D)**. In addition, we detected markers of Merkel cells, which originate from the Krt14^+^ keratinocytes (Van Keymeulen et al., 2009). Early markers such as *Atoh1, Sox2*, and *Krt8* were highly enriched in the placode population, although still expressed at low levels **(Table S1)**, a finding in line with the studies showing that Merkel cells are derived from a subset of primary hair placode cells at ∼E15 (Nguyen et al., 2019).

Of the 5 502 differentially expressed genes, 1 422 were specifically upregulated in IFE (Fc >1.5, adj p-value <0.05). Analysis of our IFE signature revealed well-established marker genes, such as *Krt1, Ovol1* and *Tgm3* **(Table S1)**. Gene sets enriched in IFE population included epidermal cell (*Tgm3, Casp14, Sprr1a*) and keratinocyte differentiation (*Tgm1, Krt10, Acer1, Foxn1, Evpl*), lipid metabolism (*Pla2g2f, Mboat2, Cds1, Lpl*) and water homeostasis (*Abca12, Aqp3, Hrnr*), as well as genes involved in calpain, IGF1 and calcineurin pathways **(Table S1 and Figure S1D)**. We also noticed high enrichment of melanoblast markers and genes involved in pigmentation and pigment and melanocyte differentiation, such as *Mitf, Kit, Sox10, Dct* and *Tyr* **(Table S1 and Figure S1D)**. This was expected, given that MAdCAM-1 is not expressed in melanoblasts.

Two studies on transcriptomes of hair follicle progenitors/IFE have been previously published. In an RNA-seq study, the entire E14.5 back skin was profiled using double-transgenic reporter mice and antibody staining; six cell populations were isolated, including P-cadherin–enriched placode cells (Sennett et al., 2015). Another, microarray-based study profiled epithelial cells responsive to transcription factor NF-κB, the main mediator of the Eda pathway (Tomann et al., 2016). Both studies confirmed our findings (**Figure S1E**). We identified 1 693 placode genes with fold-change >1.5 (adjusted p-value <0.05), whereas Tomann et al., 2016 reported 262 (Fc >1.5, adjusted p-value <0.05) out of which 253 (96.6%) overlapped with our placode marker genes (**Figure S1E, S1F and Table S2)**. Sennett et al., 2015 study defined signature genes as having at least 2-fold greater expression level in one compared to all other isolated cell types. For placode versus IFE, 1.5-fold cutoff was applied. Overlap with their placode signature was lower. Out of 102 genes, 57 overlapped with placode markers from our dataset with fold change >1.5 **(Figure S1E, S1F and Table S2)**, 22 genes had a fold change lower than 1.5 in our study, of which nine showed higher expression in IFE. Additional 21 genes were not significantly differentially expressed in our dataset **(Table S1 and S2)**.

For further validation of our hair placode signature, we chose seven genes that were upregulated in the placode population to a varying degree: *Btg1* (1.9x), *Nup210* (2.0x), *Pdzd2* (2.1x), *Alcam* (2.4x), *Stxbp6* (5.9x), *Ascl4* (11.4x), and *Cyfip2* (17.4x), and analyzed their expression by whole mount in situ hybridization (WMISH) and/or qRT-PCR at E14.5. Analysis of *Btg1* (BTG anti-proliferation factor 1), negative regulator of cell proliferation (Rouault et al., 1992), which to our knowledge has not been previously reported as placode marker, confirmed that its expression was enriched in the hair placodes **(Figure 1D and 1E)**. WMISH allowed us not only to confirm placode expression also for the other genes, and verify them as hair placode markers, but in addition suggested existence of differences in the expression area. For example, *Nup210* (nucleoporin 210), nuclear pore protein implicated in gene expression regulation during cell fate determination (D’Angelo et al., 2012) was detected in a broad area. In contrast, expression of *Alcam* (activated leukocyte cell adhesion molecule), protein involved in diverse cellular processes (Ferragut et al., 2021), with reported expression in developing whisker HFs (Fraboulet et al., 2000), and *Stxbp6* (syntaxin binding protein 6 (a.k.a. amisyn), a poorly studied protein implicated in exocytosis via regulation of the SNARE complex (Liu et al., 2021), appeared restricted to a more central area of the placode. Even more constricted central expression, similar to *Shh* (Levy et al., 2005; Heath et al., 2009), was observed for transcription factor *Ascl4* (achaete-scute family bHLH transcription factor 4), previously observed in primary hair placodes (Tomann et al., 2016), and *Cyfip2* (cytoplasmic FMR1-interacting protein 2), a gene involved in cell migration and regulation of actin polymerization through WAVE regulatory complex (WRC) (Eden et al., 2002; Derivery and Gautreau, 2010). *Pdzd2*, (PDZ Domain Containing 2), a multi-PDZ-domain protein (Yeung et al., 2003), showed a ring-like expression pattern (**Figure 1D**), a pattern previously described for *Sostdc1* (Närhi et al., 2008).

To further investigate similarities between *Shh* and *Cyfip2* expression, we used the *Cyfip2-LacZ* knockin allele as a surrogate to visualize *Cyfip2* expression in embryonic dorsolateral skin (Kumar et al., 2013) **(Figure S1G)**. Expression pattern of *Cyfip2* closely resembled that of Shh; co-localization with Shh-GFP (Harfe et al., 2004) was also detected at E15.5 **(Figure 1F)**, and similar to *Shh* (Laurikkala et al., 2002), Cyfip2-LacZ expression was absent in *Eda* null embryos and enlarged in K14-*Eda* embryos **(Figure S1H)**. Association of *Cyfip2* with actin polymerization, known to be essential for hair placode formation and polarity (Ahtiainen et al., 2014; Cetera et al., 2018), and its co-localization with Shh, led us to investigate its function in the hair placode more closely, using the *Cyfip2* null mice (*Cyfip2*^*LacZ/LacZ*^) that only survive until birth (Kumar et al., 2013). However, placode formation was undisturbed in *Cyfip2* null embryos, with normal expression pattern of *Shh* and *Dkk4* at E14.5, and the localization of Shh-GFP at the tip of the invading hair germ was unaffected at E15.5 (**Figure S1I**). No obvious morphological defects were detected at the later developmental stages either **(Figure S1I)**. This lack of mutant *Cyfip2* KO hair phenotype might be due to functional redundancy with *Cyfip1*, paralogue of *Cyfip2* (Abekhoukh and Bardoni, 2014), that was uniformly expressed within the placode and IFE **(Table S1)**. Intriguingly, a recent study suggested Cyfip2-containing WAVE complexes impair, rather than promote cell migration (Polesskaya et al., 2021).

### Single-cell transcriptional profiling of E14.5 epithelium corroborates bulk RNA-seq findings and reveals new levels of basal and suprabasal cell heterogeneity

To gain a deeper understanding of the different cell populations present in E14.5 skin epithelium, we next turned to scRNA-seq and first sequenced the entire epithelial compartment. To generate single-cell transcriptome libraries using 10x genomics Chromium system, the epithelium was processed into single cells as for bulk RNA-seq, followed by removal of dead cells using FACS. Libraries were sequenced to a depth of approximately 300 000 reads per cell, leading to identification of 4 486 genes per cell **(Figure 1A)**. After filtering, 1 601 cells were kept for further analyses. Using unsupervised clustering and t-distributed stochastic neighbor embedding dimension reduction (tSNE) we defined nine different epithelial populations and their marker genes **(Figure 2A and Table S3)**. Placode population was identified by high expression of hair placode markers *Dkk4, Edar* and *Fgf20* **(Figure 2B and 2C)**. Basal and suprabasal populations were readily identified using basal markers *Krt5* and *Krt15* and suprabasal markers *Krt1* and *Krt10*, and the melanoblast cluster by high expression of well-established melanoblast markers, including *Mitf, Sox10, Pax3, Dct* and *Tyr* (Vandamme and Berx, 2019) **(Figure 2B and 2C)**. In addition to well-established regulators, such as Mitf and Sox10, gene regulatory network (GRN) analysis by SCENIC (Aibar et al., 2017) suggests involvement of novel GRNs in melanoblast fate and/or function, including Foxo1 and Tcf12 **(Figure S2A)**.

**Figure 2.**
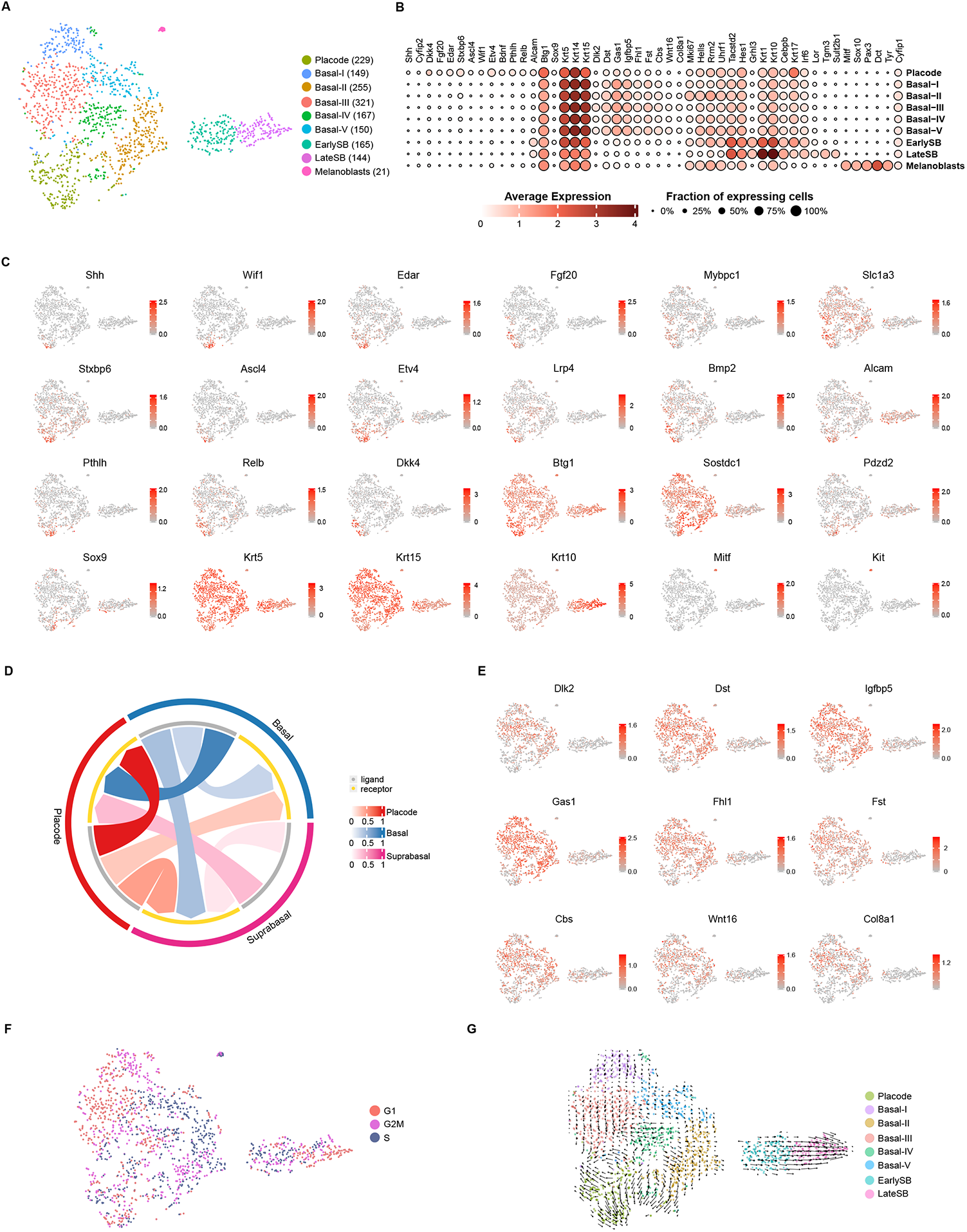
Cellular heterogeneity in E14.5 mouse back skin epithelium. (A) tSNE plot of E14.5 mouse back skin epithelium, with annotated clusters. Cell number per cluster shown in brackets. (B) Dot plot showing marker gene expression versus annotated clusters. Size of the dots represents the fraction of cells expressing the transcript, and color the average expression level within a cluster. (C) tSNE plots showing expression of cluster marker genes transcripts. Color intensity from gray to red shows the expression levels for each gene. (D) Circos plot showing global intercellular communication between cell populations. (E) tSNE plots showing expression of unique IFE basal signature genes. Color intensity from gray to red shows the expression levels for each gene. (F) tSNE plots showing cell cycle status. (G) RNA velocity field projected on the tSNE plot of mouse back skin epithelial cells. Arrows represent the average RNA velocity.

Placode cluster showed unique expression of many well-known and novel hair placode markers (*Dkk4, Fgf20, Edar, Stxbp6, Btg1*) and enriched expression of transcription factors **(Figure 2B, 2C and Table S3)**. However, no Merkel cell cluster could be discerned. When we compared placode cluster genes to our differentially expressed bulk placode markers, we found that 112 of the 113 cluster defining genes were also present in the bulk signature, further confirming our sorting strategy for the bulk transcriptomics and validating our findings **(Figure S2B)**. Some were, however, not exclusive to the placode cluster such as *Alcam*, which was identified also as a suprabasal marker **(Figure 2B, 2C and Table S3)**. Basal, suprabasal and melanoblast clusters matched our IFE bulk markers and revealed IFE bulk profile was substantially more defined by suprabasal than basal markers **(Figure S2C)**, something we could also observe in IFE signatures of previously published datasets (Sennett et al., 2015; Tomann et al., 2016) **(Figure S2D and S2E)**. Most suprabasal cluster markers overlapped with bulk IFE signature. Genes marking basal clusters overlapped in similar numbers with both placode-enriched and IFE bulk signature, with majority of them not being differentially expressed in the bulk RNAseq dataset **(Table S3 and Figure S2B)**.

### Basal IFE heterogeneity is largely cell cycle driven

Majority of analyzed IFE cells belonged to basal population, divided into five separate clusters by unsupervised clustering (Basal I-V) **(Figure 2A)**. Even though we did not observe any striking gene expression differences between IFE basal clusters, certain degree of heterogeneity, beyond cell cycle, was present. IFE basal cells (Basal I-V) expressed a number of ligands and receptors in their gene signatures and strong intercellular communication was apparent between IFE basal cells and the placode, as well as suprabasal cells, albeit to a somewhat lesser extent (**Figure 2D and S2F**). Additionally, by comparing gene expression of placode, basal and suprabasal clusters we identified nine genes, *Dlk2, Dst, Igfbp5, Gas1, Fhl1, Fst, Cbs, Wnt16* and *Col8a1*, that were expressed at significantly higher levels in IFE basal, than in suprabasal IFE or placode cells, revealing a unique IFE basal signature **(Figure 2B and 2E)**.

Despite cell cycle regression, its effect was still apparent in IFE basal clustering and cluster markers **(Figure 2F and Table S3)**. Gene ontology (GO) analysis by g:Profiler (Raudvere et al., 2019) for biological process showed enrichment of terms for cell cycle, cell division and DNA replication in three of the basal clusters (Basal-I, Basal-II, Basal-V) **(Figure S2G)**, suggesting these clusters are largely cell cycle driven, though we cannot exclude the possibility that there could be also distinct (cell cycle-independent) cell states. Namely, Basal-II and Basal-V gene signatures additionally included the highest expression of *Hells, Rrm2* and *Uhrf1* among all IFE basal cells **(Figure 2B)**. In a recent study on human neonatal foreskin epidermis, these genes were reported to mark “transitional” basal cells seemingly in the process of delaminating from the basal layer, with a possible role as basal stem cells (SC) with a fluid cell fate (Wang et al., 2020).

In contrast, Basal-III and Basal-IV did not show enrichment of cell cycle genes. Basal-III had the best overlap with IFE bulk signature **(Figure S2B)** and was defined by markers connected to HF and epidermal morphogenesis **(Figure S2G)**. Based on gene expression profile, and majority of cells being in G_1_ state, Basal-III could be considered as basal “ground” state. Basal-IV, on the other hand, could be a cluster of metabolically active basal cells, as GO analysis revealed enrichment of terms connected to metabolic processes **(Figure S2G)**.

To analyze differentiation trajectory of IFE basal cells we performed RNA Velocity, method that distinguishes between unspliced (nascent) and spliced (mature) mRNA to predict the future states of individual cells (La Manno et al., 2018). Velocity vectors from Basal-I, -IV and -V suggested differentiation trajectory going towards the “ground” state, Basal-III cluster **(Figure 2G)**.

### Differentiation trajectories reveal early suprabasal markers

Next, we had a closer look at epidermal differentiation. Unsupervised clustering identified two suprabasal clusters characterized by 487 genes in total. Majority (78%) of the markers of the first suprabasal cluster, EarlySB, were shared with LateSB cluster including many genes with established function in keratinocyte biology (e.g. *Pvrl4, Zfp750, Grhl1, Rhov, Dsc3, Ovol1, Cebpa, Cebpb*), but also several unique ones (see below) (**Figure 3A and S3A**). EarlySB still expressed a number of basal markers like *Krt5, Krt14* and *Krt15*, albeit at lower levels than basal IFE cells **(Figure 2B)**. Downregulation of the ‘genuine’ basal markers was readily evident, as was upregulation of suprabasal markers such as *Tacstd2, Krt1*, and *Krt10*. Expression profile of LateSB cluster was clearly distinct from others, with further downregulation of basal markers and upregulation of more mature differentiation markers, such as *Tgm3* and *Lor* **(Figure 2B, 3A and S3A)**. Our analysis did not reveal a specific periderm population. However, many of the established periderm markers such as *Krt17, Irf6*, and *Grhl3* (McGowan and Coulombe, 1998; Ingraham et al., 2006; Yu et al., 2006) were markers of EarlySB or both EarlySB and LateSB (**Figure 3A and Table S3**). This suggests that at E14.5 the peridermal signature is very similar to the early suprabasal IFE signature and/or the relative amount of peridermal cells was too low to distinguish them from the suprabasal cells. GRN analysis indicated increase in regulon activity going from EarlySB to LateSB **(Figure S2A)**. This trajectory was clearly shown by RNA Velocity analysis, with arrows getting longer as expression of basal markers dropped and of suprabasal increased, suggesting cells are undergoing large changes in gene expression and rapid differentiation (Svensson and Pachter, 2018) **(Figure 2G)**. Short vectors, or even vectors pointing back to basal cells, were observed in cells with intermediate expression profile between basal and suprabasal states, that still had high expression of basal, but also showed increase in expression of differentiation markers. This could point to a population of highly plastic cells, still not fully committed to the specific differentiation program, state similar to basal-suprabasal transition cells observed at postnatal day 0 (Lin et al., 2020). RNA Velocity vectors from proliferative Basal-II cluster, containing putative “transitional” basal cells, pointing towards suprabasal clusters, suggested these cells might be progenitors of suprabasal cells **(Figure 2G)**. SCENIC analysis revealed increased activity of Ezh2, E2F7 and E2F8 regulons in Basal-II **(Figure S2A)**, further implying these cells might be “transitional”: Ezh2 maintains basal keratinocytes in a proliferative state and suppresses late differentiation markers (Ezhkova et al., 2009), yet E2F7/8 have been implicated in suppression of S-phase progression and initiation of differentiation (Endo-Munoz et al., 2009; Westendorp et al., 2012).

**Figure 3.**
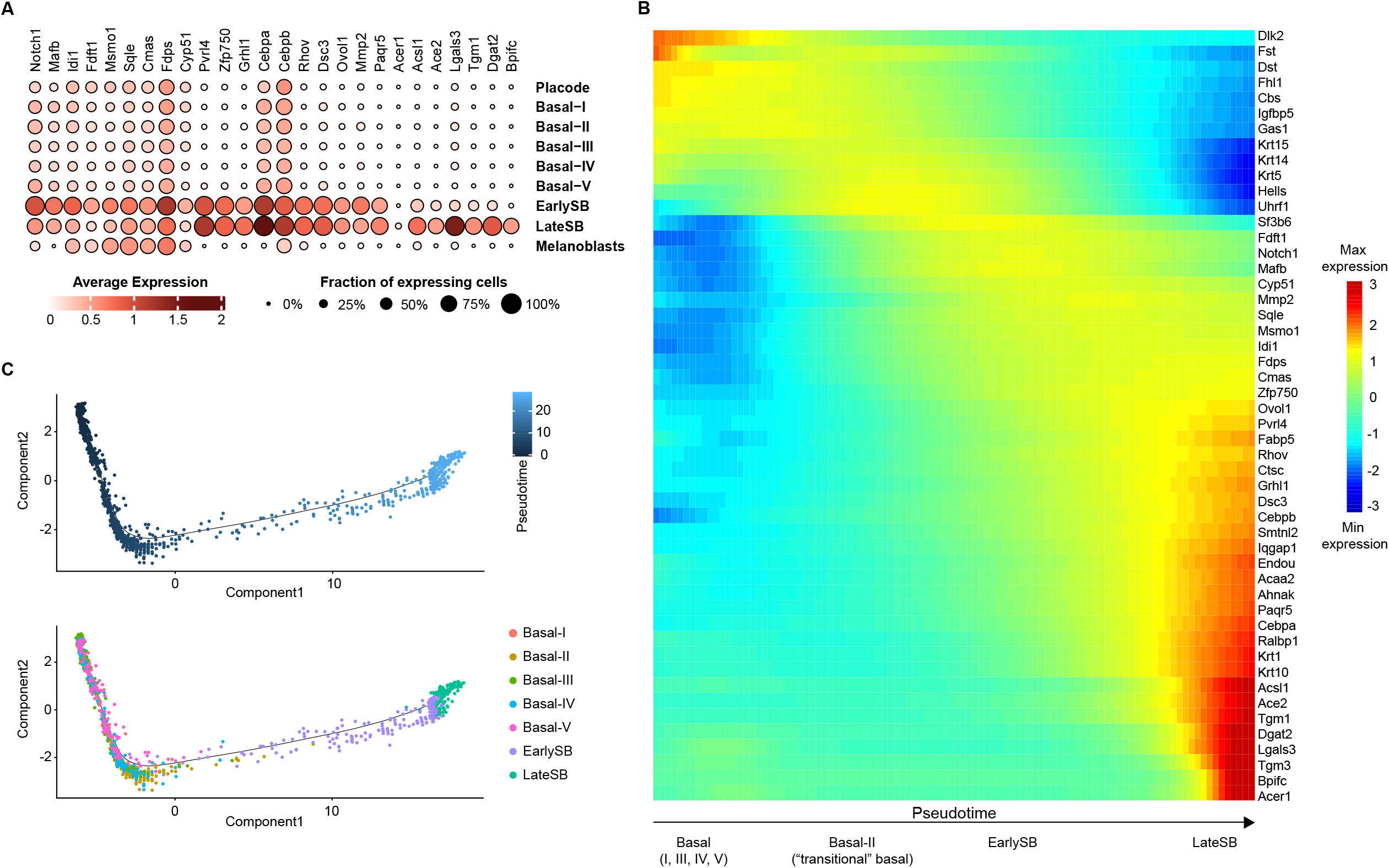
Pseudotime analysis indicates suprabasal developmental trajectory. (A) Dot plot showing expression of suprabasal markers versus annotated clusters. Size of the dots represents the fraction of cells expressing the transcript, and color the average expression level within a cluster. (B) Heat map showing expression of basal and suprabasal genes along pseudotime, for basal (Basal I-V) and suprabasal (EarlySB and LateSB) cell populations. (C) Pseudotime ordering of basal (Basal I-V) and suprabasal (EarlySB and LateSB) cell clusters.

Ordering basal and suprabasal cells along a reconstructed differentiation trajectory using unsupervised pseudotime modeling with Monocle 2 (Qiu et al., 2017) reflected RNA velocity differentiation trajectory **(Figure 3B and 3C)**. With decreasing expression of unique IFE basal markers, expression of “transitional” basal genes (*Uhrf1, Rrm2*) started to increase, until early suprabasal markers kicked in, starting with *Notch1* and *Mafb* **(Figure 3B)**, two well-known players in keratinocyte differentiation (Rangarajan et al., 2001; Miyai et al., 2016). Expression of *Notch1, Mafb*, and several other unique markers of the EarlySB such as *Idi1, Fdft1*, and *Msmo1* **(Figure 3A and S3A)**, markedly dropped with advancement of suprabasal differentiation in the LateSB **(Figure 3B)**. Temporal expression of suprabasal markers was even more evident when only EarlySB and LateSB cells were ordered along the differentiation trajectory **(Figure S3B)**. Expression of *Notch1* and *Mafb*, together with *Uhrf1, Rrm2* marked the very early stage of suprabasal differentiation, followed by a second wave of early differentiation genes, *Idi1, Fdft1, Msmo1, Sqle, Cmas, Fdps* and *Cyp51* **(Figure S3B)**. These genes have all been implicated in cholesterols synthesis (Olivier et al., 2000; Do et al., 2009; He et al., 2014; Nakamura et al., 2015; Yoshioka et al., 2020), with the exception of *Cmas*, which has an important role in formation of sialylated glycoproteins and glycolipids (Abeln et al., 2017). *Tgm1* and other late suprabasal genes started expressing towards the end of the pseudotime trajectory, along with the strong upregulation of *Acer1*, marker of differentiated epidermal keratinocytes (Houben et al., 2006), at the very of end of the trajectory **(Figure 3B, S3A and S3B)**.

### At least four cell populations define the hair placode

Even though the number of placode cells in the whole epithelium sample was relatively low, we already saw an indication of placode cell heterogeneity. RNA velocity analysis performed on all epithelial clusters, excluding melanoblasts, remarkably revealed two differentiation trajectories in the placode cluster **(Figure 2G)**. One trajectory included cells expressing *Shh, Wif1* and *Sostdc1*, while the other one was devoid of *Shh*^*+*^ and *Wif1*^*+*^ cells and mostly composed of *Dkk4*^*hi*^/*Fgf20*^*hi*^ cells **(Figure 2C)** suggesting the presence of cellular heterogeneity within the placode. To investigate this heterogeneity further we isolated a high number of MAdCAM-1-positive cells **(Figure 1A and S1A)** and generated single-cell transcriptome libraries from two independently processed biological replicates of placode-enriched cells. Libraries were sequenced to a depth of approximately 300 000 reads per cell with 4 526 genes per cell. Two samples were aggregated and after filtering 3 082 cells were kept for further analyses. Comparing computationally pooled scRNA-seq placode data to our bulk placode data confirmed the robustness of our experimental pipeline **(Figure S4A)**, a result similar to our previous comparison of methods in developing mouse teeth (Hallikas et al., 2021). Unsupervised clustering identified eight different cell populations **(Figure 4A and Table S3)**. Analysis of gene markers for each cluster revealed that MAdCAM-1-enriched population consisted of four placode clusters (Placode I-IV), three basal (IFE Basal I-III) and one suprabasal cluster (IFE SB) **(Figure 4A-C and S4B)**.

**Figure 4.**
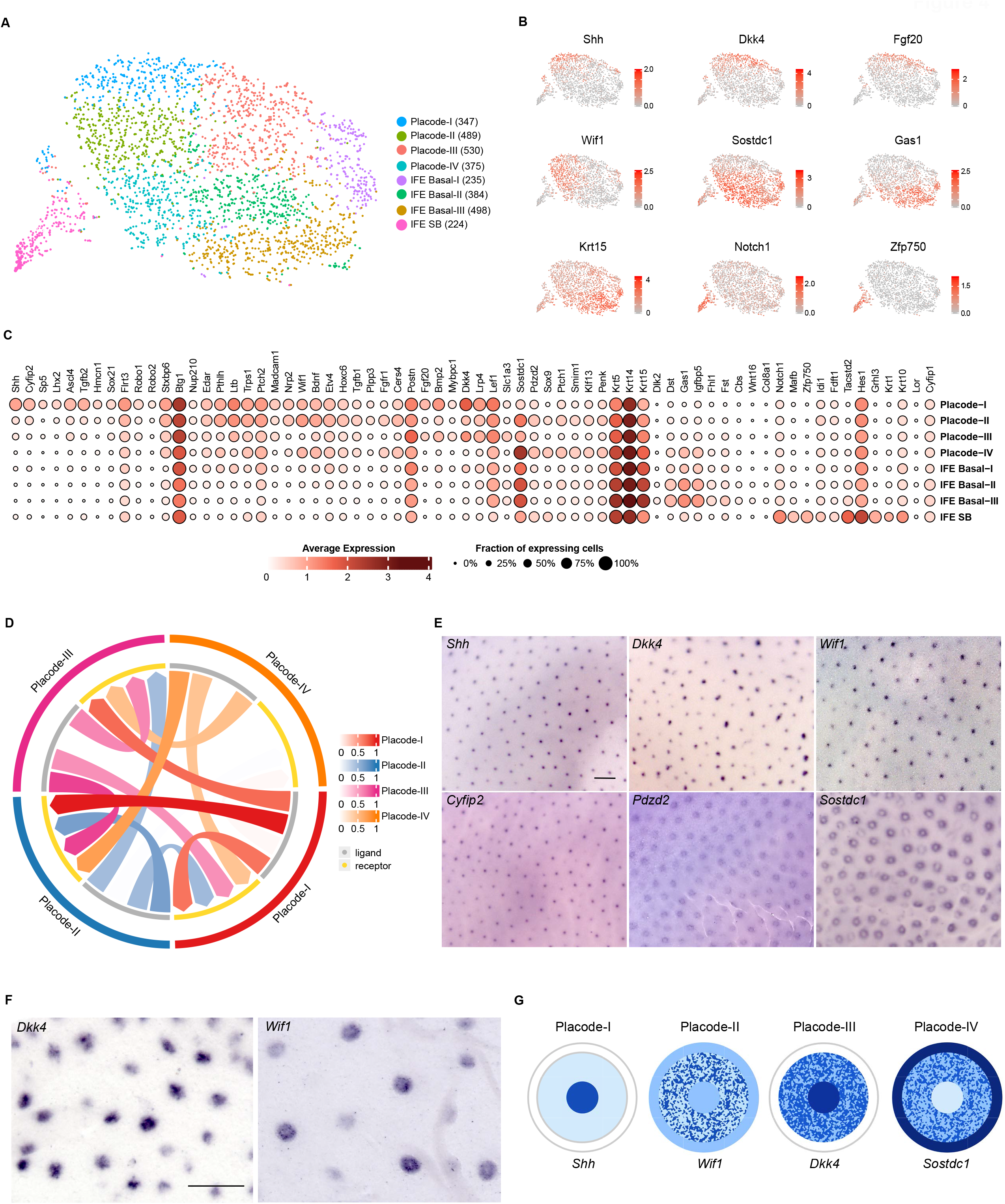
Four distinct cell populations define the hair placode. (A) tSNE plot of E14.5 hair placode-enriched population, with annotated clusters. Cell number per cluster shown in brackets. (B) tSNE plots showing expression of cluster marker genes transcripts. Color intensity from gray to red shows the expression levels for each gene. (C) Dot plot showing marker gene expression versus annotated clusters. Size of the dots represents the fraction of cells expressing the transcript, and color the average expression level within a cluster. (D) Circos plot showing global intercellular communication between four placode clusters. (E) WMISH on whole E14.5 embryos using probes specific for *Shh, Dkk4, Wif1, Cyfip2, Pdzd2* and *Sostdc1* (n = 5). Scale bar, 500 μm. (F) WMISH on E14.5 isolated back skin epithelium using probes specific for *Dkk4* and *Wif1* (n = 6). Scale bar, 200 μm. (G) Schematic of gene expression areas in the hair placode.

Each of the four placode clusters had a unique set of markers that was not shared by other clusters **(Table S4)**. Among them, Placode-I had the highest number of signature genes and unique markers, several of them involved in cell movement and migration (including *Shh, Cyfip2, Flrt3, Robo1, Robo2*) (Eden et al., 2002; Friedl and Mayor, 2017; Cetera et al., 2018; Ny et al., 2020) **(Figure 4C)**. It also had the highest number of transcription and co-transcription factors in cluster defining genes, such as the direct Wnt/β-catenin target genes *Sp5* and *Lhx2* (Zhang et al., 2008; Matos et al., 2020) **(Figure 4C, S4C and Table S3)**. Presence of *Cyfip2* and *Ascl4*, as unique markers of Placode-I cluster, together with *Shh*, further corroborated our previous observations on similarities of their expression patterns **(Table S3 and Figure 1D)**. We could also confirm the ubiquitous expression of *Cyfip2* paralogue *Cyfip1* **(Figure 4C)**, already seen at the whole epithelium level **(Figure 2B)**. Placode-II was enriched for genes connected to cell movement and growth **(Figure S4D)** and was defined by *Edar* and its target genes *Nfkbia, Pthlh, Ltb, Trps1*, and *Ptch2*, as well as *Wif1* (**Table S3**), also a gene downstream of Edar (Lefebvre et al., 2012; Tomann et al., 2016). These, and many other cluster defining genes were shared with Placode-I (**Table S4**), including *Stxbp6*, explaining its slightly broader expression pattern, compared to *Shh* (**Figure 1D**). Something we could also see for *Btg1*, shared between Placode-I and Placode-III **(Figure 1D and 4C)**. In addition, Placode-II showed high level of intercellular communication with all other placode clusters **(Figure 4D and S4E)**. Placode-III also shared a large proportion of its markers with Placode-I, but intriguingly, entirely different ones from those marking Placode-II (**Table S4**). These included many genes involved in and/or downstream Wnt signaling in HFs such as *Lef1, Dkk4, Lrp4, Fgf20*, and *Bmp2* (Zhang et al., 2008; Ahn et al., 2013; Huh et al., 2013; Matos et al., 2020). *Slc1a3*, a marker of proliferative stem and progenitor cells in adult epidermis and HF (Reichenbach et al., 2018) as well as *Mybpc1* and *Postn* were unique markers of Placode-III **(Figure 4C and Table S4)**. Finally, Placode-IV, cluster with the lowest level of intercluster communication **(Figure 4D and S4E)**, was defined by genes showing a ring-like expression, *Sostdc1* (Närhi et al., 2008) and novel placode marker *Pdzd2* **(Figure 4C and 1D-E)**, as well as early stem cell marker *Sox9* (Vidal et al., 2005; Morita et al., 2021). *Sox9*, along with most other Placode-IV markers was low or absent in Placode-I, supporting the conclusion that the *Shh*^hi^, and the prospective stem cell populations, are mutually exclusive not only in the hair germ (Xu et al., 2015; Ouspenskaia et al., 2016), but already at the placode stage, as recently shown in whisker follicles (Morita et al., 2021). Many of Placode-IV markers were unique (**Table S4**), including *Ptch1*, the commonly used readout of Shh signaling activity (**Figure S4E**), in line with notion that Shh regulates *Sox9* expression (Vidal et al., 2005; Matos et al., 2020). Another cell population high in *Sox9* expression was suprabasal cells (IFE SB) (**Figure 4C and S4B**). Two Sox9^+^ placode populations, one of prospective stem cells and another one consisting of suprabasal placode cells have recently been described (Morita et al., 2021). The IFE SB population, however, clearly does not represent a fifth, suprabasal placode population as it was devoid of all well-established placode markers including *Edar, Dkk4, Fgf20*, and instead was high in genuine suprabasal IFE markers such as *Zfp750, Notch1*, and *Mafb* (**Figure 4B, 4C and Table S3** (Placode.markers); compare to **Figure 3A, S3A and Table S3** (Epithelium.markers)). We did note, though, expression of *Sox9* in Placode-II cluster in addition to Placode-IV (**Figure 4C and S4B**).

### Hair placode populations are spatially distinct

Cluster marker analysis further strengthened our findings on gene expression differences in the hair placode. Majority of all *Shh*-expressing cells were confined to Placode-I **(Table S3; Figure 4B and 4C)** and we have already confirmed *Shh* population, marking Placode-I, was located in the center of the placode, as suggested by previous studies (Levy et al., 2005; Heath et al., 2009), together with *Cyfip2* **(Figure 1D)**. To investigate potential spatial heterogeneity in more detail, we used in situ hybridization to localize selected marker genes of other three placode clusters in E14.5 embryos **(Figure 4E)**.

*Dkk4* and *Fgf20*, markers of Placode-I and Placode-III, showed a broad expression radius, seemingly covering the whole placode area **(Figure 1B and 4E-F)**, while *Wif1* (Placode-I, II, IV), *Sostdc1* (Placode-IV; IFE Basal-II and Basal-III) and *Pdzd2* (Placode-IV unique) showed ring-like expression patterns of different diameters and placode area coverage **(Figure 4E and 4F)**. *Wif1* ring-like expression was spanning a broader placode area, starting closer to the placode center, while *Sostdc1* and *Pdzd2* rings were marking the edge of the hair placode. Because of high expression of *Wif1* in dermal fibroblast, and in particular DC (Sennett et al., 2015), its ring-like expression in epidermis was visible only after performing ISH on isolated epithelium, in the absence of mesenchyme **(Figure 4F)**.

Taking scRNAseq and ISH data together, we posit that Placode-I cells, with high expression of *Shh, Cyfip2*, and many unique marker genes such as *Tgfb2, Sox21, and Hmcn1*, is forming the placode center, and Placode-IV cells an outer ring around the whole placode, characterized by high expression of *Sostdc1, Pdzd2* and *Sox9* **(Figure 4E and 4G)**. Between these two areas we detected *Wif1*^*hi*^ zone (Placode-II), marked by Edar target genes. *Wif1* is a secreted Wnt inhibitor (Hsieh et al., 1999) and Wif1 protein has been localized to the apical side of the basal cells of the hair germ (Matos et al., 2020). Placode-I and Placode-IV also expressed *Wif1*, at somewhat lower levels than Placode-II. Interestingly, Placode-III seemed to be devoid of *Wif1* expression, but instead was characterized by expression of another Wnt inhibitor, *Dkk4*, which was in turn present at low levels in *Wif1*^*hi*^ Placode-II **(Figure 4B and S4B)**. While outside of placode center boundaries expression of *Shh* and *Cyfip2* drops sharply, *Dkk4* and *Bmp2* expression was relatively even, though slightly lower, throughout the whole placode area **(Figure 4B, 4C and 4E)**. These data suggest the area between placode center and the outer ring consists of two cells populations, one (Placode-III) defined by still rather high Wnt signaling activity. Fine gene expression gradients between placode populations were reflected in GRN analysis where regulon activity gradient was clearly visible, increasing from the placode edge towards the center **(Figure S4F)** implying regionalization may be occurring already at this early stage of hair development, with cells acquiring their fates. Lack of clear developmental trajectories between placode clusters **(Figure S4G)** was further suggestive of cell fates already being established at this stage.

## DISCUSSION

With this study, we aimed to produce a comprehensive transcriptomic profile of HF progenitors and to examine cellular heterogeneity and differentiation in HF placode and interfollicular epithelium by means of bulk and single-cell RNAseq. Utilizing physical separation of the epithelium from the underlying mesenchyme, followed by immunostaining for the cell surface marker MAdCAM-1 to isolate HF progenitors from IFE cells, proved to be a successful strategy, without the need for transgenic mice expressing artificial reporters or complex antibody panels. Analysis of bulk sequenced MAdCAM-1 positive cells showed they encompass the entirety of hair placode population, resulting in exhaustive placode and IFE gene signatures. Based on differential expression levels we were able to identify and validate several new hair placode markers such as *Stxbp6, Btg1*, and *Cyfip2*. Many of the findings from previously published datasets for early HF development (Sennett et al., 2015; Tomann et al., 2016) could be confirmed by our data, while presenting a wealth of additional insight into gene signatures of developing HFs. Expectedly, RNA-seq profiling revealed a broader gene signature than microarray approach (Tomann et al., 2016). Somewhat lower overlap with the RNA-seq study (Sennett et al., 2015) could be explained by differences in placode cell sorting strategy, number of biological replicates used, and criteria used for defining signature genes. As a result, many placode markers identified in this study had gone previously unrecognized, while some placode signature genes did not score significant in our study, as they were highly expressed also in IFE, mostly in the basal cell population. The appreciable sequencing depth obtained by bulk RNA-seq also allowed us to identify the first signs of Merkel cell specification already at E14.5, although Merkel cells could not be identified by the less sensitive scRNAseq.

Combining bulk RNA-seq with WMISH, qRT-PCR, whole mount 3D confocal imaging and literature search, gave us a glance into cell populations of the mouse back skin epithelium and the cell populations therein, but only after applying scRNA-seq we could detect the extensive cellular heterogeneity of the epithelial compartment. Bulk IFE transcriptional profile has been until now largely described by suprabasal markers (Sennett et al., 2015; Tomann et al., 2016). In contrast, the whole epithelium scRNA-seq allowed us to identify a unique IFE basal gene signature (i.e. genes expressed at higher levels in basal cells compared to both suprabasal IFE and hair placode cells) comprised of nine genes, some of them with established functions in epidermal development including *Dst, Fst*, and *Wnt16* (Guo et al., 1995; Matzuk et al., 1995; Mendoza-Reinoso and Beverdam, 2018). Additionally, a novel basal marker delta-like 2 (*Dlk2*) is a known *Notch1* inhibitor (Sánchez-Solana et al., 2011). In the early stages of epidermal development Notch signaling is required for commitment of basal keratinocytes to terminally differentiate (Blanpain et al., 2006), suggesting *Dlk2* could have a role in the maintenance of basal identity. Further studies are needed to clarify whether *Dlk2* or other IFE basal genes play any role in epidermal development.

Cellular mechanisms regulating decision between self-renewal and differentiation of basal cells are still under investigation. In the delamination model, proliferation in basal cell layer leads to cell crowding and cell-shape distortion, resulting in changes in cell-cell adhesion and cortical tension triggering differentiation, detachment from the basement membrane and upward movement (Lechler and Fuchs, 2007; Miroshnikova et al., 2018; Damen et al., 2021). In the perpendicular division model, differentiation is causally linked with the cell cycle phase and division angle: the mitotic spindle is apico-basally oriented positioning one daughter cell directly to the suprabasal layer where it begins to differentiate (Poulson and Lechler, 2010; Box et al., 2019). Our data indicate there is a population of proliferating basal IFE cells (clusters Basal-II and Basal-V) expressing markers suggestive of entering differentiation and progressing towards suprabasal fate. In a recent study on human neonatal foreskin epidermis this expression profile was described as “transitional” basal cells, purportedly in the process of delaminating from the basal membrane (Wang et al., 2020). In addition, RNA Velocity revealed differentiation trajectory going from Basal-II towards still proliferating EarlySB, further strengthening the hypothesis that suprabasal progenitors could be found in Basal-II cell cluster. Ezh2 regulon activity and high expression of *Mki67* indicate Basal-II cells are highly proliferative, yet equally distributed in G2M and S phases, a finding that does not support the perpendicular division model of differentiation. Basal-V, on the other hand, showed much lower *Mki67* expression and velocity vectors leading away from suprabasal and towards “ground” basal fate (Basal-III).

Suprabasal cells were readily distinguished from the rest of the epithelial cell populations by unsupervised clustering, having unique transcriptomic signatures separating them into two groups of early and late differentiating cells. While most suprabasal marker were shared between the EarlySB and LateSB, we also identified several early markers (*Idi1, Fdft1, Msmo1, Sqle, Fdps* and *Cyp51*), whose expression was downregulated with progression of suprabasal differentiation. Implication of many early suprabasal markers in cholesterol synthesis (He et al., 2011; Coman et al., 2018; Nagaraja et al., 2020) suggests increase in cholesterol synthesis as one of the earliest signs of keratinocyte differentiation. Cholesterol sulfate, shown to induce involucrin expression (Strott and Higashi, 2003) and act as transcriptional activator of *Tgm1* (Kawabe et al., 1998), is a metabolite of *Sult2b1*, sulfotransferase engaged in the synthesis and metabolism of steroids (Javitt et al., 2001) and one of the unique markers of LateSB. Increase of cholesterol sulfotransferase expression has been shown to accompany keratinocyte differentiation (Jetten et al., 1989).

Single-cell sequencing of a large amount of placode cells allowed us to identify four transcriptionally and spatially distinct cell populations in the hair placode: a centrally located *Shh*^+^ (Placode-I) population, a peripheral *Sostdc1*^+^/*Sox9*^+^ (Placode-IV) population, as well as Placode-II and Placode-III. A recent scRNA-seq profiling of whisker follicles suggested the existence of only two distinct cell populations at the placode stage: a central *Shh*^hi^ population, and a peripheral *Sox9*^+^ basal population of prospective stem cells (Morita et al., 2021) that apparently match our Placode-I and Placode-IV clusters. By means of in situ hybridization we confirmed the location of *Shh* expression cells to the center of the placode, as previously suggested (Levy et al., 2005; Heath et al., 2009), and other unique markers of Placode-I including *Cyfip2* and *Ascl4* display a similar pointed expression pattern. Live imaging has revealed that the initial *Shh*^*+*^ population maintains *Shh* expression and remains in close contact with the dermal condensate/papilla during later stages of HF morphogenesis indicating the fate of these cells is fixed early on (Morita et al., 2021). Placode-I has the highest levels of *bona fide* Wnt target genes such as *Sp5, Fgf20*, and *Dkk4* (Bazzi et al., 2007; Zhang et al., 2008; Huh et al., 2013), but also numerous Edar induced genes (*Fgf20, Dkk4, Lrp4, Madcam1, Ltb, Nfkbia*) (Fliniaux et al., 2008; Lefebvre et al., 2012; Tomann et al., 2016). It is possible that these cells are the first to acquire HF fate, yet their final identity is established only once they gain sufficiently high Wnt signaling activity known to depend on Edar (Zhang et al., 2009). Indeed, similar to *Shh* (Cui et al., 2011), and unlike *Fgf20* and *Dkk4* (Fliniaux et al., 2008; Huh et al., 2013), expression of *Cyfip2* was absent in *Eda* null pre-placodes. Furthermore, *Cyfip2*-LacZ was undetectable at E15.5 in nascent secondary hair placodes, which are readily visible at that stage with markers such as *Fgf20*-LacZ (Huh et al., 2013).

A ring of cells expressing *Sostdc1, Pdzd2* and *Sox9* surrounds the hair placode (Placode-IV). *Sostdc1* expression around developing hair follicles has been shown to control placode size, but not their spacing, by establishing a border between Wnt-responsive (placode) and non-responsive (IFE) cells (Närhi et al., 2012). Placode-IV population is likely established by Bmp and Shh pathways. High Bmp activity is essential for generation of the *Sox9*^*+*^ SC compartment via *Smad1/5* (Kandyba et al., 2014) and *Sostdc1* expression (Laurikkala et al., 2003; Mou et al., 2006), whereas *Shh* activity is required at least to suppress Wnt signaling (Xu et al., 2015; Ouspenskaia et al., 2016). *Sox9*^+^ cells found in Placode-IV (and Placode-II) are clearly distinct from the IFE SB *Sox9*^+^ cells. Existence of the two functionally distinct placodal *Sox9*^*+*^ populations was reported (Morita et al., 2021), the second one consisting of suprabasal *Sox9*^*+*^ cells, that based on live cell tracking do not contribute to the prospective SC compartment (Morita et al., 2021) – our Placode-II may represent this population. On the other hand, Morita et al. described the second Sox9^+^ population as “cells with flat nucleus” suggesting that they might represent peridermal cells lying atop of the stratified placode, a population of cells likely included in our *Sox9*^+^ IFE SB population.

Recent live imaging data support a ‘telescope model’ where the fate of placode cells is determined by their position with respect to the placode center (Morita et al., 2021). Hence, cells between the center and the outer ring mainly contribute to the area between the proliferative HF matrix enclosing the dermal papilla (progeny of *Shh*^+^ cells) and the bulge SCs (progeny of peripheral *Sox9*^+^ cells). Based on our in situ data, Placode-II and Placode-III, defined by high expression of Wnt inhibitors *Wif1* and *Dkk4*, respectively, do not seem to be spatially distinct populations, but rather mutually interspersed in the area between placode center and outer ring. Whether the fate of Placode-II and Placode-III cells differs in any way is currently unknown. Placode-II was characterized by Edar target genes, but appeared low in Wnt activity. On the other hand, similar to Placode-I, Placode-III signature was defined by many Wnt pathway genes including direct transcriptional targets. These two Wnt active clusters, where Wnts and Wnt inhibitors are co-expressed, would fit the model of HF progenitors oppositely polarizing Wnt activators and inhibitors, to build local boundaries and promote morphogenesis (Matos et al., 2020). In this model, cells can preserve their own Wnt^hi^ signaling identity, by localizing Wnt inhibitors apically, and at the same time prevent Wnt^lo^ cells from responding to Wnt.

Taken together, here we present a comprehensive survey of cellular populations present at developing HF and IFE and their transcriptional profiles, complementing and expanding the existing knowledge on HF and epidermal morphogenesis. Number of samples and depth of sequencing allowed unbiased gene discovery and identification of novel markers and cell populations. We reveal new basal IFE and early suprabasal markers and propose the identity of suprabasal progenitors. Unparalleled deep look into HF placode uncovers at least three spatially distinct areas composed of at least four different cell populations forming the hair placode, suggesting early establishment of cell fates. To enable further characterization of cell types, signaling pathways, and gene functions in skin and HF development, we have generated a user-friendly online tool as a resource to mine the data.

## ACKNOWLEDGMENTS

We thank Ms. Raija Savolainen and Ms. Riikka Santalahti for excellent technical assistance, and all Mikkola and Jernvall lab members for stimulating discussions. We thank Dr. Juha Kere for inspiration, and Drs. Kere and Igor Adameyko for critical reading of the manuscript. Dr. Vinod Kumar and MSc. Toiba Mushtaq are acknowledged for contributing to WMISH experiments and Mr. Hrvoje Horbec for preparing figures 1A and 4G. We thank Dr. David Rice and his lab for support in realizing FACS experiments. The mouse studies were carried out with the support of HiLIFE Laboratory Animal Centre Core Facility, University of Helsinki. Confocal microscopy was conducted at the Light Microscopy Unit, Institute of Biotechnology, supported by HiLIFE and Biocenter Finland. The flow cytometry analysis and cell sorting was performed at the Biomedicum HiLife Flow Cytometry Unit, University of Helsinki. Bulk RNA-seq was conducted in the DNA Sequencing and Genomics Unit at the Institute of Biotechnology, University of Helsinki. Single-cell RNA-seq was performed at FIMM Single-Cell Analytics unit supported by HiLIFE and Biocenter Finland. This study was financially supported by the Academy of Finland (https://www.aka.fi/en) Center of Excellence Program (grants 272280 and 307421 to MLM, IT and JJ), Sigrid Jusélius Foundation (http://sigridjuselius.fi/en) (MLM and IT), and the HiLIFE Fellow Program (https://www.helsinki.fi/en/helsinki-institute-of-life-science) (MLM). VP acknowledges support from the Finnish Cultural Foundation (https://skr.fi/en). The funders had no role in study design, data collection and analysis, decision to publish, or preparation of the manuscript.

## AUTHOR CONTRIBUTIONS

Conceptualization: A.M.S., I.T. and M.L.M.; Methodology: A.M.S., V.P., and M.M.; Software: A.M.S., R.D.R. and Q.L.; Validation: A.M.S., V.P. and R.S.; Formal analysis: A.M.S., R.D.R., V.P., Q.L. and R.S.; Investigation: A.M.S., V.P. and R.S.; Resources: J.J., I.T. and M.L.M.; Data curation: A.M.S., R.D.R. and M.L.M.; Writing – Original Draft: A.M.S., V.P. and M.L.M.; Writing – Review & Editing: A.M.S., R.D.R., V.P., Q.L., R.S., J.J., I.T. and M.L.M.; Visualization: A.M.S., R.D.R., V.P. and Q.L.; Visualizations – online tool: R.D.R.; Supervision: J.J., I.T. and M.L.M.; Project administration: A.M.S. and M.L.M.; Funding acquisition: J.J., I.T. and M.L.M.

## STAR METHODS

### RESOURCE AVAILABILITY

#### Lead contact

Further information and requests for resources and reagents should be directed to and will be fulfilled by the lead contact, Marja L Mikkola.

#### Materials availability

This study did not generate new unique reagents.

#### Data and code availability

1. All sequencing data generated in this study have been deposited at GEO and are publicly available as of the date of publication. Accession numbers are listed in the key resources table.
2. Standard programs and packages were used for data analysis in this study, and all software resources used are indicated above and shown in the Key resources table. Interactive displays of all bulk RNA-seq and scRNA-seq data can be accessed through the website.
3. Any additional information required to reanalyze the data reported in this paper is available from the lead contact upon request.

## EXPERIMENTAL MODEL AND SUBJECT DETAILS

For sequencing experiments C57BL/6JOlaHsd mice (Envigo) were used in this study. For in situ hybridization experiments NMRI mice (Envigo) were used, unless stated otherwise. C57BL/6N-A^tm1Brd^ Cyfip2^tm1a(EUCOMM)Wtsi^/WtsiBiat mice (here referred to as Cyfip2 KO), with Cyfip2-β-galactosidase knock-in allele, were obtained from the European Mouse Mutant Archive and maintained on C57Bl/6J background. Shh^GFPCre^ (Harfe et al., 2004), Fgf20 KO (Huh et al., 2012), K14-Eda (Mustonen et al., 2003) and Eda null (Pispa et al., 1999) mice have been previously described. All mice were kept in 12h light-dark cycles and food and water were available *ad libitum*. Appearance of a vaginal plug was consider as embryonic (E) 0. Embryos were individually staged according to limb and other external morphological criteria (Martin, 1990). Embryonic data are derived from stages between E14.5 and 18.5, including both sexes. All mouse studies were approved and carried out in accordance with the guidelines of the Finnish national animal experimentation board.

## METHOD DETAILS

### Replicates

Bulk RNA-seq was performed on seven biological replicates originating from seven different litters. Whole epithelium scRNAseq was performed on one replicate and placode-enriched on two biological replicates. qRT-PCR was performed on four biological replicates. For all sequencing and qRT-PCR samples E14.5 embryos from the same litter were pooled (3-4 embryos per sample) and sampled on different days. Individual stainings and ISH experiments were performed on samples originating from at least two different litters.

### Cell suspension preparation and FACS isolation

Dorso-lateral skins of E14.5 C57BL/6JOlaHsd mouse embryos were dissected and enzymatically separated using 2.5 U/ml Dispase II (Roche, Tokyo, Japan) in 4°C, 50 min, on an orbital shaker, followed by mechanical separation of epithelium and mesenchyme. Epithelial sheets of embryos from the same litter were pooled (3-4 embryos per sample) and incubated for 10 min at RT in TrypLE Select Enzyme (Life Technologies/Gibco). Dissociated cells were resuspended in 0.5% BSA, 2mM EDTA, pH 7.2 in PBS and stained with rat anti-MAdCAM-1 (1:10, BD Pharmingen) for 10 min, followed by anti-rat Alexa Fluor 488 - conjugated antibody (1:1000, Life Technologies) for 20 min, on ice. This staining step was omitted for preparation of samples for total epithelium scRNA-seq. Stained cell suspension was passed through a cell strainer and 7-aminoactinomycin D (Molecular probes by Life Technologies) was added for dead cell identification. Live placode-enriched and IFE cells, or live total epithelium cells, were sorted on BD Influx Cell Sorter (BD Biosciences) with 100 μm nozzle. Data analysis was performed using FlowJo v10.0.8r1 software (FlowJo LLC, BD).

### RNA extraction and sequencing

Sorted cells were collected directly to TRI Reagent® RT – Liquid Samples (Molecular Research Center, Inc. Cincinnati, OH). Total RNA was extracted using Direct-zol RNA Microprep kit (Zymo Research, Irvine, CA), including DNase treatment, according to manufacturer’s instructions. For RNA-seq seven biological replicates were used. RNA quality was assessed with 2100 Bioanalyzer (Agilent, Santa Clara, CA) using Agilent RNA 6000 Pico Kit (Agilent, Santa Clara, CA) according to manufacturer’s instructions. Samples used had RIN values ≥ 9.4. cDNA libraries were prepared with Ovation SoLo RNA-seq System (NuGen/Tecan Genomics), according to manufacturer’s instructions, and sequenced with NextSeq 500 (Illumina, San Diego, CA) in HiLIFE DNA Genomics and Sequencing core facility, Helsinki, Finland.

### Bulk RNA expression analysis

RNAseq reads (84 bp) were evaluated and filtered using FastQC (Andrews, 2010), AfterQC (Chen et al., 2017) and Trimmomatic (Bolger et al., 2014). This resulted with on average 60 million reads per sample. Reads were aligned with STAR (Dobin et al., 2013) to GRCm38 (mm10/Ensembl release 79). Counts for each gene were performed by HTSeq tool (Anders et al., 2015). Results are shown without normalization of gene expression based on gene length, as it does not change the pattern of results. On average 82% of reads were uniquely mapped to the genome. Differentially expressed genes were identified by DESeq2 (Love et al., 2014) and gene set enrichment analysis was done by GSEA software (Subramanian et al., 2005).

### cDNA synthesis and RT-qPCR

Sorted cells were collected directly to TRIzol reagent (Sigma-Aldrich, St. Louis, MO) and total RNA was extracted as described above. RNA purity and concentration were quantified by NanoDrop® ND-1000 spectrophotometer (Thermo Fischer Scientific, Vilnius, Lithuania). cDNA synthesis for qRT-PCR was done with iScript® gDNA Clear cDNA Synthesis kit (Bio-Rad, Hercules, CA) according to the manufacturer’s instructions, using 100 ng of RNA as input. qRT-PCR primers were designed using the following templates: Stxbp6 (NM_144552.3), Cyfip2 (NM_001252459.1), Pdzd2 (NM_001081064.2), Btg1 (NM_001731.3), using NCBI Primer BLAST software. qRT-PCR was carried out with CFX96TM Real-Time System C1000Touch Thermal Cycler (Bio-Rad, Hercules, CA) using FAST SYBR® Green master mix (Thermo Fisher), in triplicate wells. Cycling protocol was as follows: initial denaturation 95 °C for 2 min followed by 40 cycles of denaturation 5 sec at 95 °C and annealing/extension 30 sec at 60°C. qRT-PCR data was analyzed with CFX Manager (Bio-Rad, Hercules, CA) software and fold changes were calculated using the 2−△△Ct method (Livak and Schmittgen, 2001) and normalized with Hprt (Biggs et al., 2018) and GAPDH as reference genes. Statistical analyses was performed using the GraphPad Prism 7 software and two tailed Student’s T-test. For all statistical tests, p < 0.05 level of confidence was accepted for statistical significance and actual p-values (to four decimal places) were provided in the figure legends. Significance levels were defined as *p < 0.05; **p < 0.01; ***p < 0.001; ns., not significant.

### scRNAseq library preparation and sequencing

Sorted cells were collected to PBS + 0.04% BSA and transported to FIMM Single-Cell Analytics core facility, Helsinki, where cell viability was determined using LUNA-STEM™ Automated Fluorescence Cell Counter (Logos Biosystems). Single-cell cDNA libraries were prepared using the 10X Genomics Chromium Single Cell 3’ reagent v2 chemistry in accordance with manufacturer’s recommendations. Libraries were sequenced on the HiSeq 2500 System rapid mode (Illumina, San Diego, CA).

### Single cell expression analysis

Raw data processing and analysis were performed using the 10X Genomics Cell Ranger v1.3.0 pipelines by FIMM Single-Cell Analytics core facility. The “cellranger mkfastq” was used to produce fastq files and “cellranger count” to perform alignment, filtering and UMI counting. Alignment was done against mouse genome mm10. The resultant individual count data for two placode-enriched datasets were finally aggregated with “cellranger aggr”. Transcriptomes of 1701 from whole epithelium cells, and 1629 and 1661 from two placode replicates were captured. A median of 4486 and 4526 genes, and 345 113 and 305 474 reads were captured per cell, for epithelium and aggregated placode samples, respectively. Further, the filtered feature-barcode matrix was checked for quality and normalization using R package Seurat v2.3.4 (Butler et al., 2018) in R version 3.5.2/RStudio version 3.5.2. Only cells with more than 200 genes/cell and genes expressed in at least 3 cells were considered for all the downstream analysis. For a robust set of cells for the expression level calculations, we limited the analyses to cells that had number of genes > 2000, but < 7000, with less than 5% of the transcripts being mitochondrial, leaving 1601 cells from the whole epithelium and 3082 cells for aggregated placode-enriched sample. For epithelium sample genes with dispersion >0.7, and for aggregated placode >1, were used for calculating principal components (PCs). For both datasets normalization, scaling and variable gene selection was performed using the standard workflow in Seurat package. To neutralize slight batch effects, data was scaled using Seurat, for percentage of mitochondrial genes, number of UMIs and cell cycle scores. tSNE dimension reduction was performed utilizing 12 (epithelium) and 14 (aggregated placode) PCs. Unsupervised clustering was done using Seurat’s shared nearest neighbor (SNN) algorithm employing a resolution of 1 for epithelium and 0.8 for aggregated placode sample.

### Estimation of cell cycle stage for scRNAseq data

Seurat’s “CellCycleScoring” function was used for prediction of cell cycle stage using cell cycle gene list from (Macosko et al., 2015). For this purpose, expression of G1/S and S markers was used to assign S score, and of G2/M and M to assign G2M score.

### Comparison of bulk RNA-seq and scRNA-seq data

For comparison with bulk RNAseq data, single-cell data was normalized with DESeq2 (Love et al., 2014) together with the corresponding bulk RNAseq samples, and median expression levels were plotted.

### RNA velocity

RNA velocity estimation was performed with Velocyto.R v0.6 (La Manno et al., 2018). The package documentation were followed and in function “gene.relative.velocity.estimates” these values used were kCells = 20 and fit.quantile = 0.02.

### Monocle 2

Monocle version 2.8.0 (Qiu et al., 2017) in R was used to determine pseudotime developmental trajectories of suprabasal cells following package documentation. Scaled expression of curated basal, and early and late suprabasal genes ordered according to pseudotime was visualized by plot_pseudotime_heatmap function.

### CellCall

To identify molecules involved in cell-cell communication among different cell types from scRNA-seq data, CellCall (version 0.0.0.9000) (Zhang et al., 2021), a publicly available repository of curated receptors, ligands and their interactions, was applied. Gene count matrix obtained from Seurat analysis and metadata file containing the labels of each cell have been used as input for the analysis. Circular plot presenting global cell-cell communication between cell types was produced with ViewInterCircos function within CellCall package. To produce the circular plot presenting individual interacting ligand-receptor molecules, all interaction pairs identified by CellCall have been extracted and the mean expression level and percentage of expressing cells has been examined for each gene in each cell type. The plot was produced with circlize (version 0.4.15) (Gu et al., 2014).

### GO analysis

g:Profiler (Raudvere et al., 2019) was used to find enriched GO terms, category biological process, with fold change based ordered input gene lists.

### Transcription factor analysis

To identify transcription factors in placode genes from bulk-RNA sequencing, genes with fold-change >1.5 were compared against two curated human transcription factor lists (Vaquerizas et al., 2009; Yan et al., 2013) and the AnimalTFDB 3.0 database (Hu et al., 2019), and all cluster marker genes were compared for scRNA-seq data. TF list was further manually annotated.

### SCENIC analysis

SCENIC version 1.1.2 (Aibar et al., 2017) package was used for the gene regulatory network analysis with default parameters. Regulon AUC thresholds for binary activity were manually defined. For visualization, only cell types/clusters with more than 70% regulon active cells were determined as regulon active ones. The heatmap was created with ComplexHeatmap package version 2.11.2 (Gu et al., 2016).

### Histology

For histological evaluation, E15.5 bodies and E18.5 dorso-lateral skin of Cyfip2 KO embryos were collected, fixed in 4% PFA in PBS and processed for paraffin embedding using standard protocols. Transversal sections of 5μm were stained by hematoxylin and eosin, and imaged using Axio Imager M.2 widefield microscope equipped with Plan-Neofluar 20x/0.5 objective and AxioCam HRc camera (Zeiss) using bright field microscopy.

### In situ hybridization

For whole-mount RNA in situ hybridization, E14.5 mouse embryos were fixed overnight in 4% PFA in PBS at 4°C and dehydrated using increasing concentrations of methanol in PBS (25%, 50%, 75% and 100% methanol). For in situ hybridization on isolated epithelium, E14.5 dissected back skin was treated with 2.5U/ml Dispase II enzyme (Roche, Mannheim, Germany) for 50 min to separate the epithelium and mesenchyme. Isolated epithelia were fixed to 0.1 μm nucleopore filter with cold methanol for 2 min, pre-fixed in 4% PFA in PBS overnight at 4°C and dehydrated through a series of solutions of methanol in PBS. In situ hybridization was performed as previously described (Mustonen et al., 2004). Briefly, samples were rehydrated using a decreasing series of methanol, bleached with 6% hydrogen peroxide in PBS for 1h and treated with 10 μg/ml Proteinase K (Roche, Mannheim, Germany) for 20 min (5 min for epithelia) at RT. Samples were then post-fixed with 4% PFA and 0.2% glutaraldehyde (Sigma-Aldrich, St. Louis, MO). Following digoxigenin-labelled RNA probes were used: *Dkk4* (Fliniaux et al., 2008), *Shh* (Echelard et al., 1993), *Sostdc1* (Laurikkala et al., 2003),

*Wif1, Cyfip2, Pdzd2, Btg1, Alcam, Nup210* and *Stxbp6* (this study), with 1 μg/ml RNA probe in hybridization buffer at 65°C for 12h. Excess probe was removed in stringent washes. Probes were detected with BM Purple AP substrate Precipitating Solution (Roche, Mannheim, Germany), samples fixed with 4% PFA and imaged using Lumar.V12 stereomicroscope with 1.2x objective and AxioCam ICc camera (Zeiss, Oberkochen, Germany), and Lumar.V16 stereomicroscope with 1.0x objective and AxioCam 305 Color. Images were handled using ZEN 2.3 lite software (Zeiss Oberkochen, Germany).

### X-gal staining

Mouse embryos were removed from the embryonic sacs and fixed in 2% PFA, 0.2% glutaraldehyde in PBS for 2h at 4°C. Samples were rinsed with PBS and washed 3 × 15min with 2mM MgCl2 (Merck, Darmstadt, Germany), 0.02% NP-40 (Calbiochem, San Diego, CA) in PBS at room temperature. Embryos were stained overnight at RT in the dark using 2mM MgCl2, 1 mg/ml X-Gal (Thermo Fischer Scientific, Vilnius, Lithuania), 5mM K3Fe(CN)6 (Merck, Darmstadt, Germany), 5mM K4Fe(CN)6 (Merck, Darmstadt, Germany), 0.1% NP-40 and 0.2% Na-deoxycholate (Sigma-Aldrich, St. Louis, MO) in PBS. Following the staining, samples were washed several times with PBS and post fixed with 4% PFA in PBS at room temperature for 2h. Samples were imaged using Lumar.V12 stereomicroscope with 1.2x objective and AxioCam ICc camera (Zeiss, Oberkochen, Germany) and ZEN 2.3 lite software (Zeiss Oberkochen, Germany).

### Immunostaining and whole mount-confocal microscopy

Following antibodies and reagents were used for immunostaining: rat anti-MAdCAM-1 (1:1000; BD Pharmingen), rat anti-EpCAM (CD326) (1:1000; BD Pharmingen), rabbit anti-β-galactosidase (1:500; MP Biomedicals), chicken anti-GFP (1:500; Abcam), Alexa Fluor 568 (1:500; Abcam), Alexa Fluor 647 (1:500; Invitrogen), Alexa Fluor 488 (1:500; Jackson Immunoresearch) and Hoechst-33342 (1:2000; Invitrogen). For whole-mount confocal microscopy, E15.5 embryonic back skin was dissected, spread onto 0.1μm nucleopore filters and fixed with 4% PFA in PBS for 2h at room temperature. Tissues were washed in large volumes of PBS and blocked in 5% normal donkey serum (Biowest, Riverside, MO), 5% goat serum (Life Technology, Carlsbad, CA), 0.5% BSA (Sigma-Aldrich, St. Louis, MO) and 0.1% Triton X-100 (Sigma-Aldrich, St. Louis, MO) in PBS for 1h. Blocking solution was changed directly for primary antibody incubation in blocking solution and incubated for 12-48h at 4 °C. The samples were washed several times in large volumes of PBS over 6–24h. Secondary antibody and Hoechst-33342 (Invitrogen) were incubated for 6–24h, followed by washing in PBS. Samples were imaged with an upright Leica TCS SP5 and fully motorized upright laser scanning confocal microscope Leica DM6 (Leica, Wetzlar, Germany) using either HCX PL APO 20x/0,7 Imm Corr (water, glycerol, oil) or HCX APO 63x/1.30 Corr glycerol-immersion objectives. Z-stacks of samples were processed using the Leica Application Suite (LAS) X software (Leica, Wetzlar, Germany).

